# The function of macromolecular complex of CFTR-NHERF2-LPA2 in inflammatory responses of intestinal epithelial cells

**DOI:** 10.1101/186023

**Authors:** Shanshan Kong, Weiqiang Zhang

## Abstract

CFTR is a cAMP-regulated chloride channel located in the apical surface of intestinal epithelial cells; where it forms a macromolecular complex with NHERF2 and LPA2. CFTR has been shown to play a role in the pathogenies of several types of secretory diarrheas. Inflammatory bowel disease (IBD) is a chronic condition of intestine characterized by severe inflammation and mucosal destruction, genetic analysis has shown that LPA contribute to IBD and patients of cystic fibrosis also display the phenotype of diarrhea. The purpose of this study is to investigate if this complex plays a role in the inflammatory responses of intestinal epithelium.

We then explored the role of this complex in maintaining the integrity of tight junction and inflammatory responses in these cells. In vitro assays show that inhibiting CFTR or LPA2 in the intestinal epithelial cell could disrupt the epithelial cell junction, and reduce the TER of intestinal epithelial cells in both mouse and human cell line. EƯSA assay show that intriguing LPA2 through LPS or LPA can increase the secretion of IL-8, while inhibiting or SiRNA knockdown of LPA2 can decrease the secretion of IL-8 in mouse or human intestinal epithelial cells. The CFTR inhibitor can reduce the IL-8 secretion in both mouse and human cell line, the deletion of CFTR in mouse intestine does not affect the IL-8 level, but the knockdown of CFTR in human cell line reduced the IL-8 protein level. The deletion of CFTR in human also reduced the IL-8 mRNA level. This indicates the CFTR-LPA complex is necessary for the expression of IL-8.

## Introduction

Intestinal epithelial cells (IEC) are polarized cells, the apical side faces the intestinal lumen with the apical membrane forming micro folds, and the basal side rests the basal membrane playing a role in communicating with blood stream and lymphatic lacteals. The apical and basolateral membranes are separated by tight junctions which are key for the regulation and transport of ions and liquids. IEC can form a continuous physical barrier that separate mammalian hosts from the external environment[1][2], and has diverse functions including physical segregation of commensal bacteria and the integration of microbial signals: form a barrier in the intestine to prevent the entrance of harmful substances; secrete digestive enzymes; exchange the water and electrolyte; secrete mucus and the innate immune function. IEC are crucial mediators to regulate the normal immunological function of intestinal[3]. Dysregulated epithelial barrier function may lead to inflammatory bowel disease which is associated with the increased bacterial translocation. And increasing evidences also indicate that loss of epithelial barrier function contributes to systemic immune activation, which promote the onset of immunological diseases[4]. Therefore, comprehensive understanding the immune regulatory properties of intestinal epithelial cells could aid to develop new strategies to prevent and treat human infection and inflammatory and metabolic diseases.

CFTR is a chloride channel that localized at the apical membrane of epithelial cells, and has been shown to play important role in the cystic fibrosis diseases[5]. Cystic fibrosis, also known as CF is a common disease that are inherited and mostly found in young population. It is an autosomal recessive disorder which means that a person must receive two altered CF genes in order to get this condition. It is a life-threatening disorder that causes severe damage to the lungs and digestive system. Millions of Americans, estimated ~30% Americans carry the defective CF gene, but do not show any symptom. The average life span for people with this disease is approximately 37 years, which is a much more than what it used to be. CFTR is critical for remaining the normal function of epithelial cells in fluid homeostasis, airway fluid clearance, and airway submucosal glands secretion[6]. The dysfunction of CFTR can lead to clinical symptoms affecting predominantly the lungs and the gastrointestinal system. As the main CΓ channel in the apical membrane, defective CFTR usually leads to aberrant ion and fluid homeostasis at the epithelial cell surface. Patients with this condition produces thick, sticky mucus, which clogs the lungs, causes repeated infection and difficulty breathing. In the digestive system, the thick mucus can also block tubes that carry digestive enzymes from pancreas to intestine, as a result the intestine aren’t able to absorb the nutrients without these digestive enzymes. The CFTR is also highly expressed along the intestinal tract, especially in the crypts in the small intestine and near the base of the crypts in the large intestine[7]. The CF also can lead to digestive symptoms in patients including foul-smelling, greasy stools; poor weight gain and growth and intestinal blockage and severe constipation nose. Along with the elongated life expectance, the digestive complications have increasingly become an important cause of morbidity in CF patients.

CFTR is critical for remaining the normal function of epithelial cells in fluid homeostasis, airway fluid clearance, and airway submucosal glands secretion[6]. In the small intestine Cl-secretion drags Na+ and water across the tight junctions, and the pathophysiological changes of trans-epithelial transport leading to chronic diarrhea[8]. Na^+^/H^+^ exchanger 2 and 3(NHE-2, NHE-3) located at the apical side of the epithelial cells, exchangers can be coupled with Cl^-^/HCO3^-^ exchanger. Studies has found that Na+/H+ exchanger regulatory factors regulate CFTR membrane retention, conductivity, and interactions with other transporters. The Na+/H+ exchanger regulatory factor (NHERF) family consists of four scaffolding proteins, including NHERF1, NHERF2, NHERF3, and NHERF4[9]. They contain two or four PDZ domains which serve as protein-protein interacting sequences and highly expressed in epithelial tissues. Pulmonary infection with an exaggerated inflammatory response is the major cause of morbidity and mortality in cystic fibrosis, LPS stimulation increased the inflammation cytokines. People have shown that wild-type (WT)-CFTR forms a macromolecular complex with NHERF2 and LPA2 at the apical plasma membrane of intestinal epithelial cells and airway epithelial cells[10]. CFTR form macromolecular complexes with other proteins at the plasma membrane of gut epithelia, which functionally couple LPA2 signaling to CFTR-mediated chloride transport[11], LPA2 is a G-protein-coupled transmembrane receptor for Lysophosphatidic acid (LPA), which is a bioactive phospholipid with diverse effects on various cells. LPA can inhibit CFTR-mediated chloride transport through the LPA2- mediated Gi pathway, and LPA inhibits CFTR-dependent cholera toxin-induced mouse intestinal fluid secretion in vivo[12]. These results indicate that LPA2-elicited signaling regulates CFTR. But the function of CFTR-LPA2 pathway in intestine has not been evaluated.

Intestinal epithelial cell function impairment is related with IBD. Epithelial cell function in the intestinal immunity pathway. Intestinal epithelial cell secrete basolateral IL-8 rapidly after the stimulation of bacterial, the IL-8 is a pro inflammatory interleukin which is demonstrated to be important to support neutrophil infiltration in ulcerative colitis and psoriasis. Signaling pathways that are involved in neutrophil migration across the mucosal barrier in the intestine have been defined in vitro. <u>The integrity of epithelial cell in the immunity function</u>.

In this paper we show that the CFTR-LPA2 pathway plays important role in regulating the immunity function of intestinal epithelial cells. The interaction of CFTR-NHERF2-LPA2 can be detected in the intestinal epithelial cells. And in vitro assays show that inhibiting CFTR or LPA2 in the intestinal epithelial cell could disrupt the epithelial cell junction, and reduced the TER of intestinal epithelial cells. In vivo experiment also show that intriguing LPA2 increased the expression of IL-8, while inhibiting LPA2 or SiRNA knockdown of LPA2 can decrease the expression of IL-8.show that inhibit CFTR or intrigue LPA can affect the fluid homeostasis of intestinal. Our results found that the CFTR-LPA2 pathway play important role in regulating the normal function of intestinal epithelial cells.

## Results

### Interaction of CFTR-NHERF2-LPA2 in mICC12 cells

As m-ICC12 cells are intestinal epithelial cells providing physical and biochemical barrier between commensal and pathogenic microorganisms[13], they can sense and respond to microbial stimuli to reinforce their barrier function and to participate in the coordination of appropriate immune responses, ranging from tolerance to anti-pathogen immunity. Thus, IECs maintain a fundamental immune regulatory function that influences the development and homeostasis of mucosal immune cells.

To investigate the function of CFTR-NHERF2-LPA2 complex in m-ICC12 cells, we first confirmed the interaction of the three molecules in mouse intestinal cells. The CFTR-IP results show that, the CFTR, NHERF2 and CFTR, LPA2 can form a complex, this indicated that the three molecules interact with each other in intestinal epithelial cells. The immunostaining results also demonstrated that NHERF2 and LPA2 are co-localized with CFTR. The above data show that the CFTR, NHERF2 and LPA2 form micro complex in mouse intestinal cells. We also detected the mRNA level of LPA receptors in m-ICC12 cells by Q-PCR, the results show that the expression levels of LPA2 and LPA3 were relatively high compared with other LPA receptors. This indicates LPA2 may play more crucial role in regulating the normal function of intestinal epithelial cells.

### CFTR-NHERF2-LPA2 complex regulates the integrity of tight junction in intestinal epithelial cells

The intestinal epithelium has complex mechanisms to control and regulate bacterial interactions with the mucosal surface. Apical tight junction proteins are critical in the maintenance of epithelial barrier function and control of par acellular permeability[14]. The characterization of alterations in tight junction proteins are key players in epithelial barrier function in inflammatory bowel diseases is rapidly enhancing our understanding of critical mechanisms in disease pathogenesis as well as novel therapeutic opportunities. The disruption of integrity of the epithelial barrier may intrigue the onset of inflammatory bowel diseases[15]. The leaky intestinal epithelial barrier is mainly attributed to defects of the TJs and IEC loss[14]; the deficient TJs are the primary cause for the compromised intestinal epithelial barrier[3].

IBD is a chronic condition of intestine characterized by severe inflammation and mucosal destruction. In the normal intestine, epithelial tight junctions provide barrier function to prevent the diffusion of bacterial, toxins, allergens from the gut lumen into intestinal tissue and systemic circulation. Genetic analysis has shown that LPA contribute to IBD and patients of cystic fibrosis also display the phenotype of diarrhea[16]. Drugs perturb CFTR-containing macromolecular complexes in the intestinal can disrupt the TJ structure and also affected the immune function of epithelial cells.

To detect whether CFTR-LPA2 pathway play important function in the cellular junction in the intestinal epithelial cells. We add chemicals that can disrupt or intrigue CFTR-LPA2 pathway to the epithelial cells and then using ZO-1 antibody to detect the cellular junction integrity. Results show that, ImCc12 cells treated with CFTR-inh172, C143 and LPS can disrupt the cellular junction. We also repeated this experiment in human cell line HT-29cells, the result show that, the cells that treated with CFTRinh172, C143 and LPS for 20 hours display disrupted cellular junction. This indicate that, the CFTR-LPA2 pathway play important roles keeping the cellular junction integrity in both human and mouse epithelial cells.

To get further evidence about the role of LPA2 in maintaining the integrity of epithelial cell tight junction, we measured the TER(trans epithelial resistance) of HT-29 cells after treatment with LPS (10ng/ul) or C143 (10uM and 20uM) for 24 hours. TER is a widely accepted quantitative technique to measure the integrity of tight junction dynamics in cell culture models of endothelial and epithelial monolayers, TER values are considered to be strong indicators of the integrity of the cellular barriers[17]. The results show that LPS treatment can significantly reduce the TER level of HT-29 cells, and the C143 at the concentration of 20uM can also reduce the TER level of HT-29 cells. This result display the function of LPA2 in maintain the TER of epithelial cells.

### CFTR-NHERF2-LPA2 complex regulates IL-8 expression in intestinal epithelial cells

Previously, people have found that intestinal epithelial cells (IEC) can secret IL-8 in response to challenges from such as lipopolysaccharide with butyrate and IL-1□[18]. IL-8 is a chemokine that stimulates migration of neutrophils from intravascular to interstitial sites and can directly activate neutrophils and regulate the expression of neutrophil adhesion molecules. To determine the function of CFTR-NHERF2-LPA2 in the process of IL-8 secretion, we treated mouse intestinal cell line and human colon cell line with molecules that block or intrigue CFTR-LPA2 pathway, and then detect the IL-8 in both mRNA and protein level. Q-PCR results show that, intrigue LPA2 through LPS can also upregulate the mRNA level of IL-8 both in HT-29 cells and mICc12 cells; block LPA2 through C143 decreased the mRNA level of IL-8 in HT-29 cells. This indicate the CFTR-LPA pathway play important role in regulating the IL-8 transcription. We also detected the IL-8 level through ELISA to confirm the function of CFTR-LPA pathway in regulating IL-8 level. The ELISA result displaying the similar results: intriguing LPA2 through LPS can elevate the IL-8 protein level in cell medium supernatant and block LPA2 through C143 downregulate the IL-8 protein level. These data indicate LPA2 positively regulate the expression and secreting of IL-8 of intestinal epithelial cells. The effect of LPA2 in IL-8 secretion also can be demonstrated through the LPA (Lysophosphatidic acid) activation. Since LPA can intrigue LPAR2, the treatment of LPA at conferential concentrations of 10uM to 40uM in HT-29 cells and mICC12 cells can stimulate the IL-8 concentration in cell medium.

To confirm the function of LPA2 in regulating the expression of IL-8 in epithelial cells, we reduced the LPA2 expression in HT-29 cells and mICC12 cells through RNA knock down. The western blot results show that the LPA2 expression can be successfully reduced through the transfection of LPA2 lentivirus, and the mRNA level of IL-8 is reduced when LPA2 is knocked down in both HT-29 cells and mICC12 cells. This experiment indicate that the IL-8 expression can be regulated through LPA2 receptors.

## Discussion

Dysfunction of intestinal epithelial cells are related with some kind of immunological diseases, such as diarrhea, and intestinal bowel diseases. Restoring the epithelial cell function would be a novel therapy strategy for IBD therapy. Previously people has found that, CFTR-NHERF2-LPA2 form macrocomplex in intestinal epithelial cells, inhibition of CFTR through CFTR inh-172 attenuates diarrhea in DSS-induced colitis[19]. Chemicals that inhibit CFTR-LPA2 interaction can lead to diarrhea in mouse, this indicate the function of CFTR-LPA2 in intestinal. In this paper, we demonstrated this macro complex also formed in the intestinal epithelial cells in vitro. CFTR IP-western can drag down the NHERF2 and LPA2 in mouse intestinal cells, LPA assay also demonstrate the interaction of CFTR-LPA2 in the in vitro cell lines.

In healthy epithelial cells, the apical TJs construct a dynamic intestinal barrier that regulates the paracellular uptake of water, nutrients and electrolytes. TJ dysfunction can lead to the disruption of the intestinal barrier integrity, and can lead to inflammatory bowel diseases. Barrier defects contribute to diarrhea by a leak flux mechanism (e.g., in IBD) and can cause mucosal inflammation by luminal antigen uptake, and the barrier dysfunction happens early in ulcerative colitis. Abnormalities in epithelial tight junction is a major defect of barrier function. Previous data has found that CFTR interacts with ZO-1 at tight junctions through its PDZ-binding domain and CFTR regulate tight junction assembly and control tubular genesis in cultured epididymal epithelial cells[20]. They also found that CFTRinh172 can significantly reduce the transepithelial electrical resistance. The most frequent CFTR gene mutation, F508del, has been shown to be associated with a disorganized actin cytoskeleton and altered tight junction permeability[21]. In our study, we demonstrate that CFTR is highly expressed in the intestinal epithelial cells, and the inhibition of CFTR or intriguing of LPA2 can lead to the disruption of tight junction of epithelial cells in both mouse and human. The treatment of LPS and C143 reduced the TER of intestinal epithelial cells. The impaired tight junction may contribute the increased inflammation in epithelial cells.

Intestinal epithelial cells secrete the chemokine IL-8 in response to bacterial entry and in the onset of acute of IBD, IL-8 can attract and active leucocytes, recruit PMNs to the inflamed crypt and the intestinal lumen to amplify or sustain the inflammation[22]. Previous study found that the CFTR knock down can increase the expression of IL-8 in CF airways as the imbalance between pro-inflammatory and anti-inflammatory[23]. In our present study, we demonstrated that intrigue LPA2 elevated IL-8 expression at both the mRNA and protein levels, while inhibiting or SiRNA knock down of LPA2 reduced the IL-8 level. Together, in this paper we show that CFTR-LPA2 involved in the intestinal inflammation.

---

**Fig.1.**
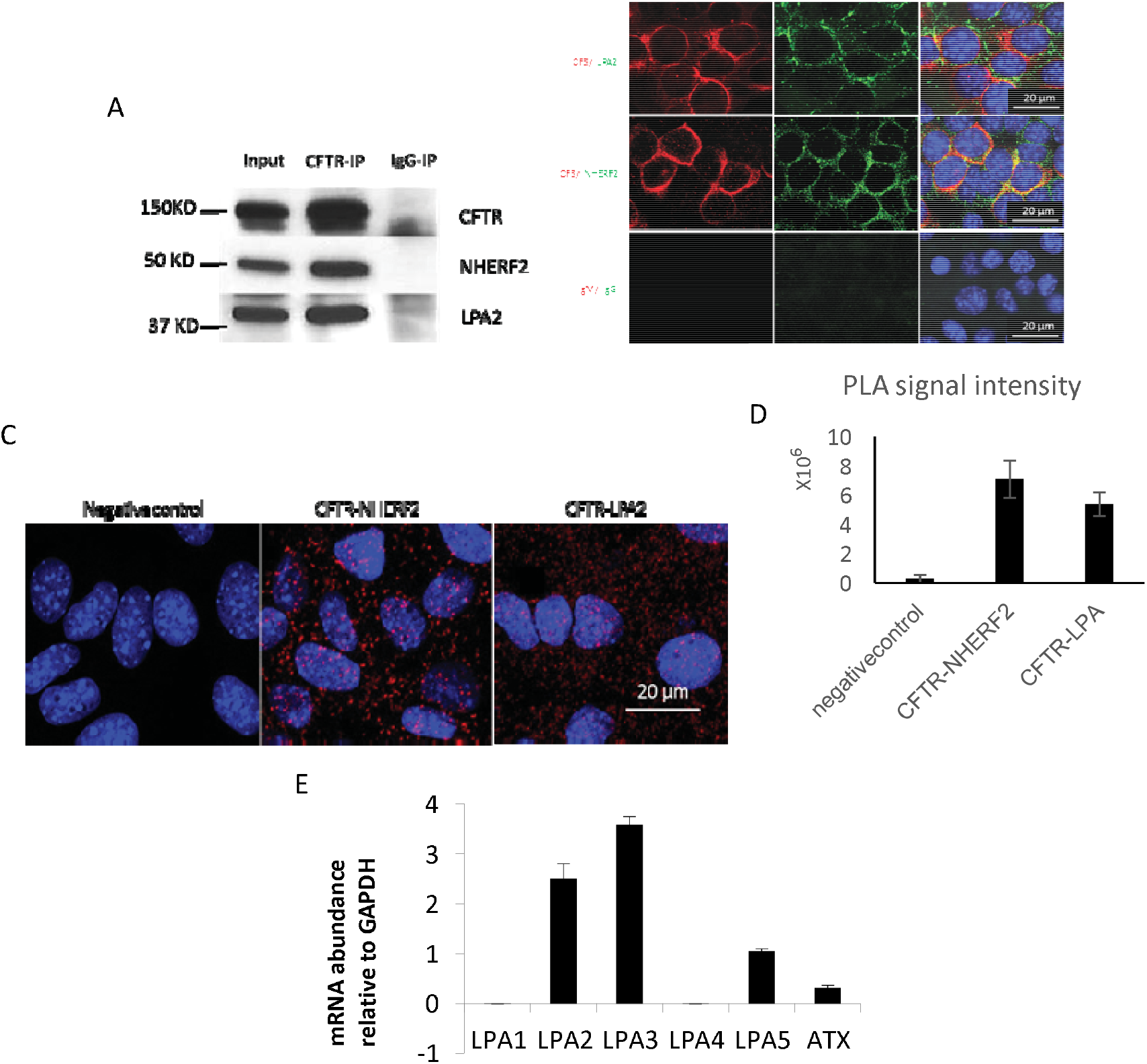
LPA2 form a complex with CFTR and is highly expressed in intestinal epithelial cells. (A) A representative blot showing that NHERF2 and LPA2 could be co-immunoprecipitated with CFTR in mICc12 cells. Mouse IgG was used as a negative control of CFTR antibody for co-IP experiments. IP: immunoprecipitation. IB: immunoblotting. (B) NHERF2 and LPA2 are co-localized with CFTR in mICc12 cells. The cells were permeabilized and stained with mouse α-CFTR and rabbit α-NHERF2 (or rabbit α-LPA2) antibodies, followed by incubation of fluorescence-labelled secondary antibodies and subjected to confocal microscopy. Mouse IgG and rabbit IgG were used as negative antibody controls. The nuclei were stained with DAPI. (C) Duolink proximity ligation assay to detect protein interactions of CFTR and NHERF2, CFTR and LPA2 in mouse intestinal cell line m-ICCl12. Duolink^®^ proximity ligation assay (PLA) showed the interactions between CFTR and NHERF2, and CFTR and LPA2 in mICc12 cells. The cells were treated with mouse α-CFTR and rabbit α-NHERF2 (or rabbit α-LPA2), followed by incubation with anti-mouse-minus probe and anti-rabbit-plus probe and followed the manufacturer’s instructions. Each red spot represents an interaction between the binding partners. The nuclei were stained with DAPI. The corresponding IgGs were used as negative controls. (D) PLA data quantification using ImageJ software. (E) Normalized mRNA expression of LPA receptors in m-ICC12 cells. Q-PCR were used to detect the gene expression level of LPA receptors (LPA1-LPA5). The results show that LPA2 and LPA3 were highly expressed in m-ICC12 cell line. The relative mRNA levels were normalized with GAPDH. Results were reported as means ± SD (error bars)

**Fig2.**
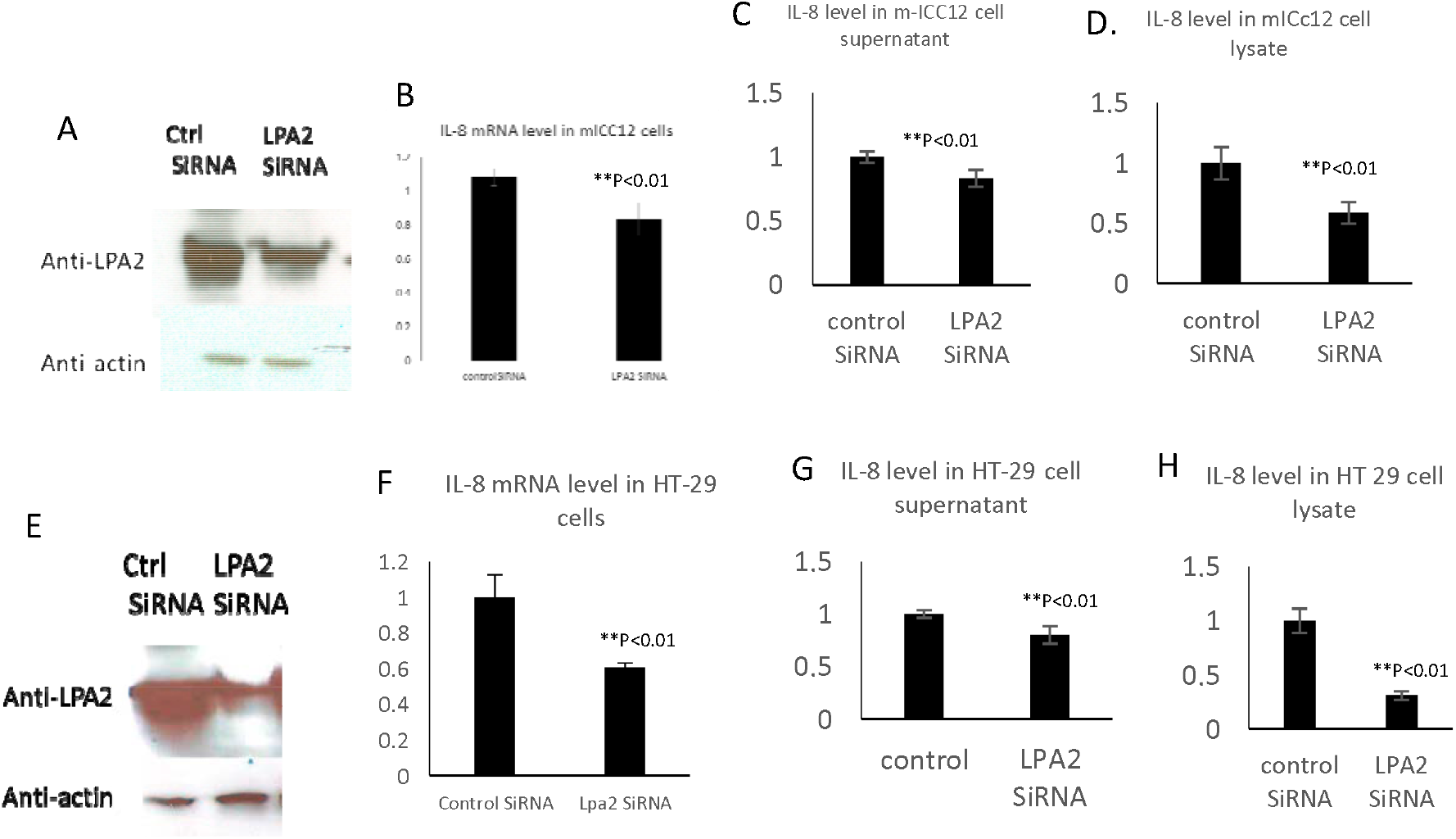
Knock down LPA2 reduced the IL-8 expression and secretion in intestinal epithelial cell. Western blot detect the LPA2 protein level in mICc12 cells (A) and HT-29 cells (E). The LPA2 expression level was reduced by RNAi. The mICC12 cells or HT-29 cells were infected by lentiviral vector carrying shRNAi against LPA2, the LPA2 expression was analyzed by western blotting. (B) The relative IL-8 mRNA levels in mICc12 cells transfected with LPA2 SírNa lentivirus and control SiRNA lentivirus. The cells were lysed with Qiagene mRNA kit to extract mRNA and Q-PCR was used to quantify IL-8 mRNA levels. (C and D) The IL-8 level in mICC12 cell supernatant or lysates were measured by ELISA, the LPA2 knock down can reduce the IL-8 level. (E) The LPA2 expression was analyzed by western blotting after the HT-29 cells were infected by lentiviral vector carrying shRNAi against LPA2. (F) The relative IL-8 mRNA levels in HT-29 cells transfected with LPA2 SiRNA lentivirus and control SiRNA lentivirus. The cells were lysed with Qiagene mRNA kit to extract mRNA and Q-PCR was used to quantify IL-8 mRNA levels. (G and H) The IL-8 level in HT-29 cell supernatant or lysates were measured by ELISA, the LPA2 knock down can reduce the IL-8 level. Each assay was carried out in triplicate, with results reported as means ± SD (error bars).*P<0.05. **P<0.01.

